# Extinction Risk from Human Impacts on Small Populations of Marine Mammals

**DOI:** 10.1101/2020.03.28.013698

**Authors:** Paul R. Wade, Elisabeth Slooten

## Abstract

Population models used to set limits for whaling, fisheries bycatch and other human-caused mortality (HCM) usually focus on relatively large populations and do not include Allee effects (declines in population growth rate at small population sizes). These models are not suitable for managing small and endangered populations of marine mammals. We use a stochastic age-structured population model to investigate the effect of HCM on extinction risk. Compared to environmental variability and catastrophes, Allee effects had a strong influence on risk. Depending on the scenario, HCM (1) delayed the rate of population recovery (with no increased risk), (2) increased extinction risk because populations lingered at low levels, (3) increased extinction risk because the population was pushed below an Allee threshold, or (4) increased extinction risk over 100 years because the rate of extinction for a doomed population was accelerated. Population dynamics in small populations are poorly known for most marine mammals. Therefore, we recommend that managers consider the range of potential population dynamics for the species under consideration and make precautionary decisions on allowable levels of HCM. For critically depleted populations (e.g., small populations, well below historic levels) even low levels of HCM have the potential to substantially increase extinction risk.

## INTRODUCTION

Small populations are vulnerable to the risk of extinction and human-caused mortality (HCM) that reduces those populations further will contribute towards this risk (Petersen and Levitan 2001). Methods developed to calculate quotas for whaling and marine mammal bycatch in fisheries (e.g., Kirkwood 1992, Cooke 1994, 1995, Wade 1998, Slooten 1998, Gambell 1999, Taylor *et al.* 2000) ensure that a relatively large population stays at or above the level at which it is most productive. However, these models do not investigate the increased threat of extinction that could be caused by HCM nor do they consider the population dynamics of very small populations. For small and endangered populations, a different modeling approach is needed that merges stochastic models used in population viability analysis (Beissinger and Westphal 1998) with models used to examine the effect of HCM on populations (e.g., Wade 1998). This paper presents a method of quantifying the increased risk of extinction caused by deliberate or unintentional removals of individuals from small populations.

The processes that cause small populations to have a greater risk of extinction include genetic and behavioural problems, as well as chance processes like demographic stochasticity (Gilpin and Soule 1986, Goodman 1987a, Lande 1993, Shaffer 1981, Simberloff 1988). A positive relationship between population size and per capita population growth is often referred to as the “Allee effect” or “depensation” (e.g., Allee 1931, Allee *et al.* 1949, Dennis 1989, 2002, Fowler and Baker 1991, Pfister and Bradbury 1996, Courchamp *et al.* 1999, Stephens and Sutherland 1999, Stephens *et al.* 1999, Petersen and Levitan 2001, Berek et al. 2007). In essence, as the number of individuals decreases there are costs from a lack of predator saturation, impaired anti-predator vigilance or defence, a breakdown of cooperative feeding, an increased possibility of inbreeding depression or other genetic issues, decreased mating due to not finding mates, or a combination of these effects. The Allee effect increases risk to small populations directly by contributing to the risk of extinction, and indirectly by decreasing the rate of recovery of exploited populations and therefore maintaining populations at a smaller size where extinction risk is higher for a variety of reasons (e.g., environmental or demographic stochasticity) (Dennis 1989, Stephens and Sutherland 1999).

It is very difficult to detect Allee effects for practical and theoretical reasons (Lande 2002), and little is known about these processes for most endangered species (Brook *et al.* 1997, Ralls *et al.* 2002). Likewise, the exact nature of the stochastic elements that affect a population’s dynamics are rarely known. Ralls *et al.* (2002) recommend incorporating several alternative forms of model dynamics in population viability analyses (PVA). Therefore, we chose to examine the effect of HCM across a wide variety of different forms of stochastic dynamics, including combinations of different types at the same time. The aim is to explore all feasible scenarios (Beissinger and Westphal 1998) and examine whether there are any common results that emerge to guide us in the face of uncertainty about any particular population’s dynamics.

Our case study is a “generic” marine mammal species, with a life history similar to dolphins. However, the approach is flexible and can be applied to marine mammals with different life history characteristics. It would also be useful for assessing the impact of intentional and accidental mortalities on a range of other species, such as kills of terrestrial animals crossing roads (Clevenger et al. 2003; Malo et al. 2004) or kills of birds in wind farms (Johnson et al. 2002, Barrios and Rodríguez 2004).

Wade (1998) investigated the effect of HCM on marine mammal populations and proposed a mortality limit (*ML_Rec_*) that could be used for species given special protection status because they are at a low level. This limit was designed to achieve a specified goal for the rate at which a severely depleted population recovers; for example, find the level of mortality that does not delay by more than 10% the time required for a population to return to 50% of carrying capacity in a deterministic model. However, managing HCM for endangered species involves some special considerations, because any HCM may be important to populations at extremely low abundance. Currently no quantitative method exists for objectively evaluating what level of HCM, if any, can be viewed as acceptable in managing endangered species of marine mammals, even though such evaluations are required under, for example, the U.S. Endangered Species Act and Marine Mammal Protection Act. The effect of HCM needs to be evaluated in the context of how much it might increase the risk of extinction for the population, which was not considered in the investigations of Wade (1998). In other words, a PVA (Gilpin and Soule 1986) that considers factors such as environmental and demographic stochasticity is necessary for investigating the effect of HCM on small populations.

Our model incorporates the standard aspects of stochastic dynamics: demographic variance, environmental variance and catastrophic mortality (e.g., Paine 1979, Shaffer 1981, Beissinger 1986, Shaffer 1987, Ewens *et al.* 1987, Goodman 1987a, 1987b, Simberloff 1988, Pimm 1991, Dennis *et al.* 1991, Lande 1993, Gaston and McArdle 1994, Mangel and Tier 1994, Beissinger 1995, Ludwig 1999, White 2000). We use an age-structured population dynamics model including demographic variance in births, natural deaths, and human-caused deaths, auto-correlated environmental variance in survival, and random mortality events that represent catastrophes. The model features “regular” density dependence (compensation) as well as a flexible function for Allee effects that is scaled (through a threshold parameter) to a biologically relevant reference point – the population size at which the model switches from positive to negative expected population growth.

Our inability to reliably determine the population dynamics of small populations is a major difficulty in evaluating extinction risk for real populations. Primarily for this reason, several authors have cautioned against the use of PVA models to estimate absolute extinction risk, but have instead advocated for the use of PVA to evaluate and compare conservation and management options (Beissinger and Westphal 1998, Morris and Doak 2002, Ralls et al. 2002). We propose evaluating risk in marine mammals from HCM by comparing extinction risk in scenarios with and without HCM. The quantity of interest is the level of HCM that increases the risk of extinction (relative to a no-HCM scenario) by a specified threshold. This level is compared across many scenarios to determine if a consistent pattern emerges. The goal is to determine an acceptable level of HCM that will be robust to our lack of knowledge of the expected population dynamics of a specific real population.

## METHODS

### Population model

The basic model was a simplified age-structured model with different survival rates for the first age class (yearlings), juveniles, and adults. After the first year, the juvenile survival rate applied until the age of first reproduction for both males and females. All mature females had the same fecundity rate until they reach maximum age. In contrast to a typical Leslie matrix model, the model was explicitly integer-based, meaning the number of individuals alive in any age class was always an integer. All births and deaths were added or subtracted as “whole” individuals (details below).

Compensation density-dependence (where population growth rates slow as a population approaches “carrying capacity”) is generally thought to be an important feature of large-mammal population dynamics (Fowler 1981, 1987), and can have some implications for extinction risk (Sabo et al. 2004). Compensation density-dependence on survival of all age classes beyond the first age class was incorporated using a generalized logistic function:

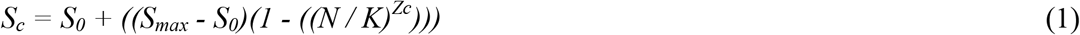

where,

*K* = carrying capacity
*N* = current population size
*S_c_* = survival rate for the given population size, under compensation
*S_0_* = survival rate resulting in zero population growth
*S_max_* = maximum survival rate
*Z_c_* = shape parameter that determines whether compensation response is linear (*Z_c_*=1) or convex (*Z_c_*>1).

A maximum value (*S_max_*) was set for both adult and juvenile survival, and they were both scaled proportionally by the same amount according to Equation 1. Further modifications to the survival rate (see below) applied to both adult and juvenile survival. This represents an age-structured version of the generalized logistic model used in Wade (1998).

The following function was used to model the Allee effect (positive relationship between population size and population growth rate) at low population sizes:

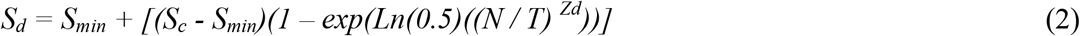

where,

*S_d_* = survival rate influenced by compensation and depensation, depending on the given population size
*S_min_* = minimum survival rate
*T* = threshold population size where expected population growth is zero (stable) under Allee effect
*Z_d_* = shape parameter that controls the steepness of depensation (Allee effect)
*S_min_* was determined by iteratively solving for the survival rate (*S_0_*) that results in zero expected population growth, and then letting

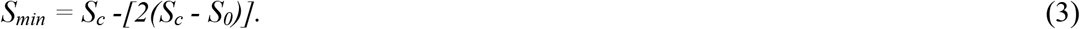

When *S_min_* was calculated in this way (where *S_min_* is the same distance below *S_0_* as *S_max_* is above *S_0_*) the survival rate will be *S_0_* when the population size equals *T* (and therefore the expected population growth will be zero). Equation 2 models an Allee effect in a similar fashion as the two-parameter model described by Stephens and Sutherland (1999), but this formulation allows explicit control of the threshold where negative population growth is expected, as well as the steepness of the effect. Figure 1 compares two versions of this Allee model (with *Z_d_*=1.5 and *Z_d_*=10.0) to a density-dependent model with no Allee effect, where *T*=100.

**Figure 1.**
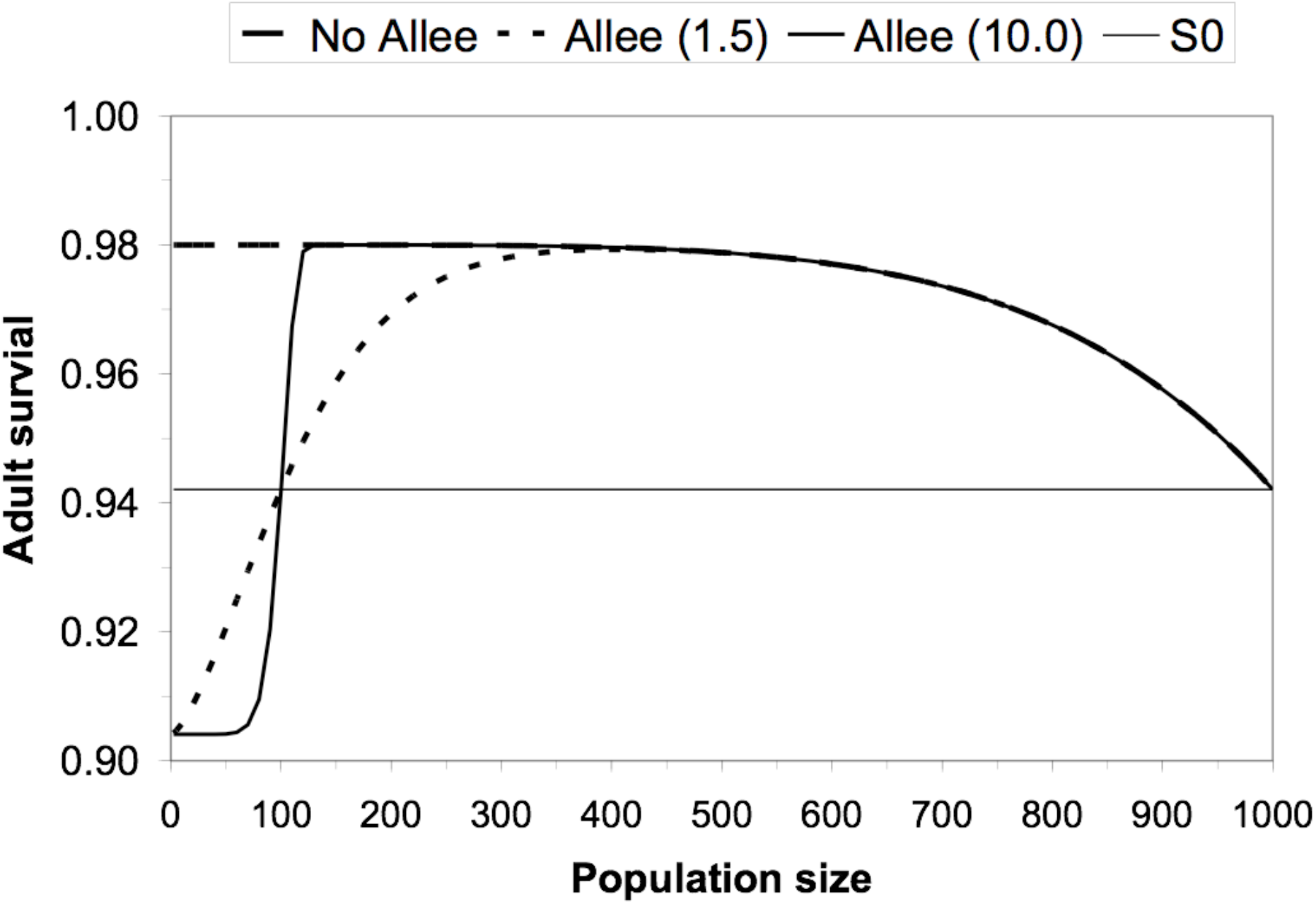
Density-dependent adult survival function used in the population model. “No Allee” is the function with no Allee effect, “Allee (1.5)” is the Allee effect with *Z_d_* = 1.5, and “Allee (10.0)” is the Allee effect with *Z_d_* = 10.0. The adult survival level (*S_0_*) that results in zero population growth (λ=1.00) is also plotted for reference.

Environmental variance was incorporated by sampling survival rates from a Beta distribution with a mean equal to the expected survival rate (as determined by Eq. 1 and 2) and a standard deviation determined by *SD_env_*. The Beta distribution is particularly useful because it is bounded between 0.0 and 1.0, and a sample of Beta random deviates can be generated that will have the exact mean and standard deviation desired. In the sequence of the model, the environmental variance calculation follows the density-dependence calculation described above, and therefore modified *S_d_*, the survival rate that takes into account density-dependent compensation and depensation.

The following function allows auto-correlation to be incorporated into environmental variance, leading to series of consecutive years with relatively good survival and relatively bad survival rate:

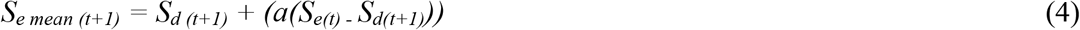

where,

*a* = an auto-correlation parameter, which takes a value between 0 (no correlation) and 1 (complete correlation)
*S_d (t+1)_* = density-dependent survival rate for year *t+1*
*S_e (t)_* = survival rate subject to (density-dependence and) environmental variability, in year *t*
*S_e mean (t+1)_* = the expected (mean) survival rate for year t+1 incorporating environmental variability.

The realized survival rate in a given year, *S_e (t+1)_*, was a random deviate from a Beta distribution with mean equal to *S_e mean (t+1)_* and a variance specified as *SD_env_* squared. Each random survival rate in a given year was therefore potentially a function of the realized survival rate the year before. Note that *SD_env_* represents the standard deviation of the Beta distribution around *S_e mean (t+1)_*, but does not represents the standard deviation of survival from year to year, which increases with values of a > 0.0.

When *a* = 0, there is no auto-correlation. In that case, a random deviate was sampled from a Beta distribution centered on the expected survival rate (*S_d (t+1)_*) that is calculated by the two density-dependent equations (Eq. 1 and 2). When *a*=1, a random deviate was sampled from a Beta distribution centered on the survival rate that was experienced in the previous year (*S_e (t)_*). This represents a random walk process, where survival is purely a function of survival in the previous year, and thus removes the effect of the density-dependent functions. When *a* is some value between 0 and 1, the effect is to create a mixture of these two extremes. For example, when *a* = 0.5, a random deviate is sampled from a Beta distribution centered on the point exactly half way between the expected survival rate from the density-dependence equations and the survival rate experienced in the previous year. This will cause the model to respond more slowly to density-dependent processes and have runs of years with bad survival and with good survival.

Mortality events (catastrophes such as disease outbreaks or sources of HCM that are not “direct” such as oil spills) were incorporated using two parameters, the probability of a mortality event (*P_event_*) and the magnitude of an event (*M_event_*). A uniform random deviate was used to determine if an event occurs. In a year in which a mortality event occurs, the survival rate was further modified as:

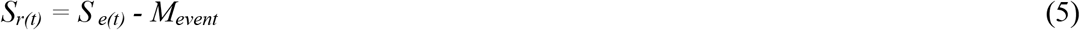

where *S_r(t)_* was the realized survival rate in year *t* after the effect of density-dependence, environmental variance, and catastrophes.

Finally, the realized survival rate *S_r(t)_* occured as a stochastic process to incorporate demographic variance, through the use of a binomial distribution. The number of random deaths in an age class were determined by taking the number of individuals in an age class as the number of trials and sampling a number of deaths from a binomial distribution with probability equal to 1.0 minus the realized survival rate for that age class in that year. Similarly, random births were determined by taking the number of mature females as the number of trials, and sampling a number of births from a binomial distribution with probability equal to the fecundity rate. This is appropriate for a life history where females only have one offspring at a time (i.e., no twinning). Sampling births and deaths in this way is identical to determining individual fates one at a time, as in an individual-based model, but was more efficient in terms of computation time.

The population was initialized by calculating the stable age distribution associated with the Leslie matrix defined by the survival and fecundity rates, where the survival rates included the effects of Eq. 1 and 2 (but not the effects of environmental variance or catastrophes). Then the starting population size was randomly distributed to age classes according to the proportions of the stable age distribution, where the number of individuals in each age class was a random draw from a binomial distribution with probability equal to stable age distribution proportion in that age class, with the number of trials equal to the starting population size (i.e., a random sample from a multinomial distribution).

### Human-caused mortality

Mortality from human sources was modeled where the expected HCM was a constant fixed proportion of population size. This represents a constant risk model of future mortality, where every individual has a constant risk of being killed from human causes. One example of this type of mortality model would be bycatch of a marine mammal in a fishery that had constant effort through time (over years). Under this model, the number of individuals killed in each year will decline or increase with population size, but the expected proportion of the population killed will stay constant. This source of mortality was subtracted from the realized survival rate calculated in Eq. 5, so that there was demographic variance in the HCM as well as in the other sources of mortality. Each scenario was run for eight different levels of HCM (0.000, 0.00125, 0.0025, 0.005, 0.010, 0.020, 0.030, and 0.040).

In summary, the flow of events in the model is as follows.

1. For each new trajectory, calculate the stable age distribution associated with the starting population growth rate, and distribute individuals among age classes,
2. For each year in a particular trajectory, calculate the survival rate associated with the population size last year, incorporating both compensatory and depensatory density dependence,
3. Use that survival rate as the basis for a distribution from which to randomly sample a survival rate to represent environmental variability (incorporating auto-correlation with last year’s realized survival if specified),
4. Randomly determine if a catastrophe occurs, and if so, subtract that mortality from the randomly sampled survival rate,
5. Subtract the specified amount of Human Caused Mortality from the survival rate.
6. Use this final, realized survival rate to determine the number of deaths in a (binomial) stochastic process,
7. Determine the number of births in a similar binomial process.

### Parameter values (common to all analyses)

The life history used for our analysis was designed to reflect a typical dolphin, a relatively long-lived mammal, with a relatively old age of sexual maturity, and a relatively low maximum rate of increase (Table 1). A single life history was used to investigate the effect of different levels of HCM and different assumptions about potential impacts to the population. It is well known that small populations of cetaceans are subject to HCM. For example, the North Atlantic right whale (~300 individuals) is subject to HCM from ship collisions and fishing gear entanglement (Kraus 1990). Maximum adult survival was set at 0.98, maximum juvenile survival at 0.95, and maximum yearling survival at 0.832. The mean birth interval was set at 3 years, so the fecundity rate (births per female) was set at 0.33. Age of first reproduction was set at 7 years, and maximum age was set at 30 years, with females reproducing up to age 30. In combination, these values of life history parameters will yield a λ of 1.04, a growth rate thought to be typical for a dolphin (Reilly and Barlow 1986). The highest level of HCM investigated (0.04) was therefore equal to the maximum population growth rate possible.

**Table 1.** Summary of parameter values fixed over all scenarios.

| Parameter | Value |
| --- | --- |
| $S_{max}$ for adult survival | 0.98 |
| $S_{max}$ for juvenile survival | 0.95 |
| Yearling survival | 0.832 |
| Mean birth interval | 3 |
| Age at first reproduction | 7 |
| Age at last reproduction | 30 |
| Maximum age | 30 |
| Carrying capacity ($K$) | 1000 |
| Allee threshold ($T$) | 100 |
| Density-dependent shape parameter ($Z_c$) | 5 |
| Number of years to project | 100 |
| Number of trajectories | 5000 |

Cetaceans are thought to have density-dependent population dynamics (Fowler 1984). The shape parameter for density-dependence (*Z_c_*) was set to a value of 5, which results in a non-linear function, as suggested for marine mammals (Fowler 1987, 1994, Taylor and DeMaster 1993). It has also been suggested that marine mammal population dynamics are influenced by Allee effects (Fowler and Baker 1991) and catastrophes (Harwood and Hall 1990, Young 1994, Gerber and Hilborn 2002). The threshold (*T*) for the Allee effect was set at 100. Other parameter values specified to describe the full population dynamics are described below in the description of different scenarios.

Five different starting population sizes were used: 50, 75, 100, 125, or 150 individuals. These sizes were chosen to span a range from below to above the Allee effect threshold. It should be noted that the role of demographic variability will change for different population levels, so our results will not necessarily be universal for all small populations, as the specific results will be conditional on these starting assumptions. A constant carrying capacity was specified of 1000. In this case, carrying capacity was sufficiently greater than the starting population size that compensation density-dependence would have little or no effect on the starting population and would serve only to prevent simulated populations from expanding without limit once they obtained population sizes greater than about 500.

Each scenario was projected 100 years and repeated 5000 times. Extinctions were tallied when the population fell below a size of two individuals.

### Population Dynamics Scenarios

The dynamics of small populations are poorly determined. Therefore, a variety of different scenarios were run with different stochastic effects (Table 2). A standard model was specified that incorporated only demographic variance and a relatively small amount of environmental variance (with no auto-correlation), with *SD_env_*=0.01. Analyses were run adding to the standard model three different forms of stochastic dynamics: (1) additional environmental variance, (2) an Allee effect, and (3) catastrophes. Scenarios were run by turning on, in turn, each of the three forms of stochastic dynamics, with both a base level and a greater level of the effect. Finally, two combinations of all three components were run.

**Table 2.** Specification of parameter values for all scenarios.

| Scenario | Allee<br>effect<br>($Z_d$) | Auto-correlation<br>($a$) | Environmental<br>variance<br>($SD_{env}$) | Probability of<br>catastrophe<br>($P_{event}$) | Severity of<br>catastrophe<br>($M_{event}$) |
| --- | --- | --- | --- | --- | --- |
| 1: Environmental Variance | 0.00 | 0.00 | 0.04 | 0.00 | 0.00 |
| 2. Auto-correlated Env. Variance | 0.00 | 0.80 | 0.04 | 0.00 | 0.00 |
| 3. Catastrophes (0.02, 0.25) | 0.00 | 0.00 | 0.01 | 0.02 | 0.25 |
| 4. Catastrophes (0.03, 0.40) | 0.00 | 0.00 | 0.01 | 0.03 | 0.40 |
| 5. Allee (1.5) | 1.50 | 0.00 | 0.01 | 0.00 | 0.00 |
| 6. Allee (10.0) | 10.00 | 0.00 | 0.01 | 0.00 | 0.00 |
| 7. Combination A | 1.50 | 0.00 | 0.04 | 0.02 | 0.25 |
| 8. Combination B | 10.00 | 0.80 | 0.04 | 0.03 | 0.40 |

The first level of environmental variance, with no auto-correlation, was set at *SD_env_* = 0.04. For example, with this level of variability and an expected survival rate of 0.80 the realized survival rate would fall within the range 0.72-0.88 with 95% probability. Note that with the relatively high adult and juvenile survival rates specified here and with the use of the Beta distribution, the distribution of realized survival rates would not be symmetric around the expected value (due to being constrained by an upper bound of 1.0), but would still result in the arithmetic mean of the random survival rates being equal to the expected value. For the scenario with a greater level of environmental variance, *SD_env_* was again set at 0.04, but the auto-correlation (*a*) was set to 0.8, which would yield a strong correlation in survival rates from year to year, mimicking sequences of years with poor or good conditions. Note that the auto-correlation increases the realized variance in survival over what is specified in the parameter SDenv; in this specific case the standard deviation of environmental deviations was ~0.06. Therefore, this scenario both increases the variance and causes it to be auto-correlated.

The first level of catastrophes was set at an expected frequency of two catastrophes in 100 years, with a magnitude of 0.25. This means that in the year of a “catastrophe”, the value of was subtracted from the expected survival rate (e.g., adult survival rate of 0.98 would become 0.73). The second catastrophe level was set at an expected frequency of 3 catastrophes every 100 years, with a magnitude of 0.40.

The first Allee effect was set at *Z_d_* = 1.5, and the second was set at *Z_d_* = 10.0. Under the first scenario the Allee effect occurs gradually, starting well above the specified threshold, and not reaching full effect until well below the threshold (Fig. 1). Under the second sceanario the Allee effect occurs steeply at a population level close to the threshold, and reaches full effect just below the threshold.

For each starting population size within each population dynamics scenario, the level of HCM was found that increased by 0.1 the proportion of trajectories that went extinct compared to the same scenario run with no HCM. These threshold levels were then plotted against starting population size. In some cases, no such value could be found. First, in a scenario where the proportion of trajectories that went extinct with no HCM was >0.90 it is not possible to increase extinction risk by 0.1. Second, in some scenarios the extinction proportion never exceeded 0.1 even at the highest level of HCM. In those cases, no value was plotted.

## RESULTS

For most scenarios, but not all, extinction risk in 100 years was very low when HCM was zero. As expected, the probability of extinction was higher for small starting population sizes and high levels of HCM. All scenarios had some extinction risk at the highest levels of HCM.

### Environmental variance (no auto-correlation)

Environmental variance has relatively little effect on extinction probability (Fig. 2A), especially at low levels of HCM. Trajectories increase relatively rapidly towards carrying capacity over 100 years, and therefore “escape” extinction risk (Fig. 3A). With HCM at 0.02, some trajectories are initially prevented from increasing (Fig. 3B), because the expected growth rate has been lowered by HCM increasing the probability of populations staying at low initial numbers for many years. At 0.04 HCM, populations tend to decline towards extinction. However, at low population density (and maximum growth rates) trajectories nearly follow a random walk, with a nearly equal probability of increasing or decreasing and some trajectories going extinct within 100 years.

**Figure 2.**
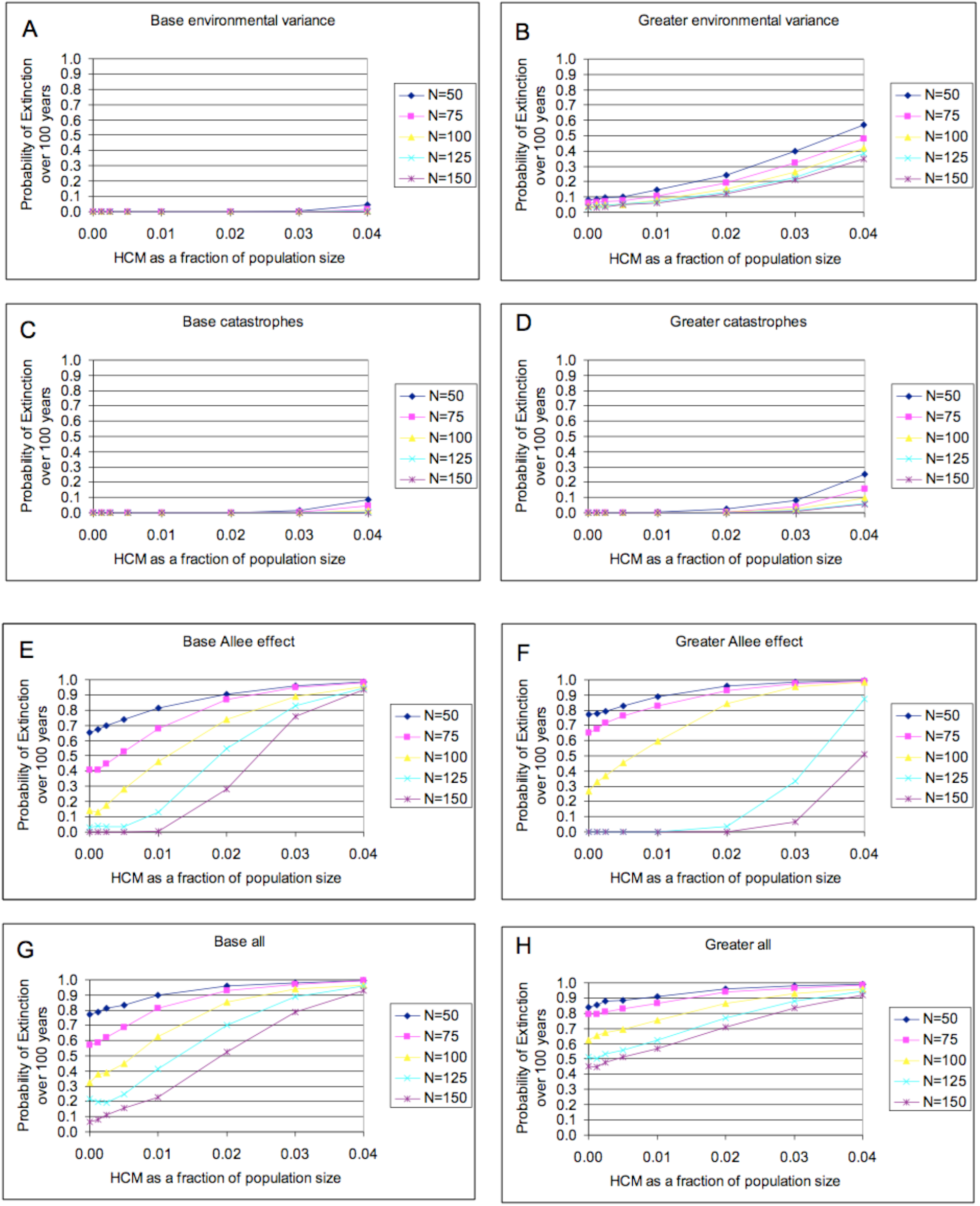
Probability of extinction for (A) Scenario 1: Environmental Variance, (B) Scenario 2: Auto-correlated Environmental Variance, (C) Scenario 3: Catastrophes (0.02, 0.25), and (D) Scenario 4: Catastrophes (0.03, 0.40), (E) Scenario 5: Allee (1.5), (F) Scenario 6: Allee (10.0), (G) Scenario 7: Combination A, and (H) Scenario 8: Combination B. See Table 2 for a specification of each Scenario. In each panel, results from simulations for 5 different starting population sizes (from 50 to 150 individuals) and for 8 different levels of human-caused mortality are shown.

**Figure 3.**
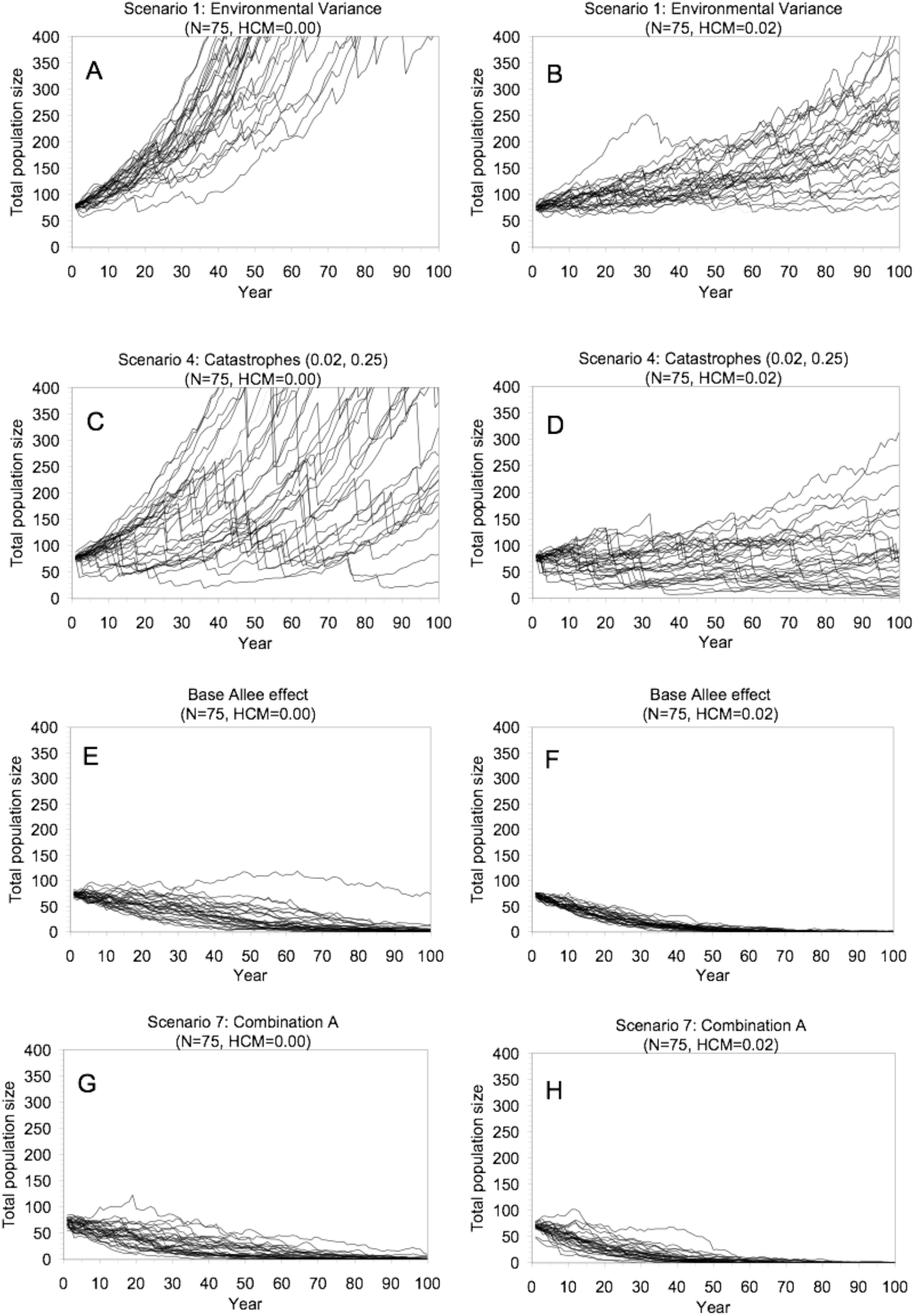
Example trajectories illustrating two types of small population response to HCM. Shown are 30 randomly chosen trajectories from the 5000 that were run. The first type of response is where HCM delays the rate of recovery of a population but does not increase extinction risk – (A) Scenario 1 with a starting population size of 75 with no human-caused mortality, versus (B) Scenario 1 with a starting population size of 75 with a human-caused mortality rate of 0.02. The second type of response is where HCM causes a population that would otherwise likely recover to linger at low population levels, increasing its risk of extinction from stochastic processes – (C) Scenario 4 with a starting population size of 125 with no human-caused mortality, versus (D) Scenario 4 with a starting population size of 125 with a human-caused mortality rate of 0.02. The third type of response is where HCM causes a population that would otherwise likely recover to have a greater probability of falling below an Allee threshold (if there is one), and thus decline to extinction – (E) Scenario 6 with a starting population size of 75 with no human-caused mortality, versus (F) Scenario 6 with a starting population size of 75 with a human-caused mortality rate of 0.02. The fourth type of response is where HCM accelerates the rate of extinction for a population that is already doomed to extinction, if current conditions continue – (G) Scenario 7 with a starting population size of 125 with no human-caused mortality, versus (H) Scenario 7 with a starting population size of 125 with a human-caused mortality rate of 0.02.

### Greater environmental variance (with auto-correlation)

This scenario produces a small chance of extinction with no HCM, for all starting population sizes (Fig. 2B). Extinction risk increases greater than linearly with HCM, and a lower starting population size raises the entire line. For example, with an initial population size of 75, there is a probability of extinction of 0.058 with no HCM and 0.107 with a HCM rate of 0.01. Correlated variance in population growth rate results in long periods of increase or decline, which can lead to extinction risk (Fig. 3B). Higher starting population sizes result in lower extinction risk (Fig. 2B). Adding HCM lowers the rate at which trajectories “escape” from low levels, and therefore increases extinction risk because there is a greater probability a population will stay at low levels where extinction can occur if it experiences a sequence of “bad” environmental years.

### Base catastrophe level

With no HCM, there was essentially no risk of extinction for populations with the base catastrophe frequency and magnitude we specified (Fig. 2C). Because the effect of catastrophes is expressed as a decline in the survival rate, their magnitude in terms of the number of individuals lost to the population becomes less when a population is small. The effect of “catastrophe” events was not large enough to overcome the overall positive growth rate of the density-dependent model (Fig. 3C). At 0.02 HCM some trajectories stayed at low population levels (Fig. 3D).

### Greater catastrophe level

With a greater frequency and magnitude of catastrophic events there was still no extinction risk for populations with no HCM (Fig. 2D). Extinction risk did occur at 0.02 HCM or greater. With the greater magnitude of catastrophe events, trajectories can quickly drop to low levels. With HCM at 0.02 and a starting population size of 75 many trajectories gradually decline towards extinction, even though few extinctions occurred by year 100 (Fig. 3D). Populations that started at a level of 125 individuals have a lower probability of going extinct, but some of these trajectories stay at population levels below 100 for the entire 100 years.

### Base (gradual) Allee effect

The base Allee effect (*Z_d_* = 1.5) resulted in relatively large changes in extinction risk. Scenarios with small starting populations, below the Allee threshold (set here at 100) had extinction probabilities in the region of 0.40-0.70, even without HCM (Fig. 2E). Populations starting at or above the Allee threshold had a much lower extinction risk (0.00 to 0.13). Populations starting below the threshold are prevented by the Allee effect from recovering (Fig. 3E), and HCM hastens the extinction of what are essentially “doomed” trajectories (Fig. 3F). Even for a population starting at 150, well above the threshold of 100, the probability of extinction is 0.28 with HCM at 0.02, which represents an initial mortality from HCM of 3 individuals per year.

### Greater (steeper) Allee effect

The steeper Allee effect (*Z_d_*= 10) led to greater extinction risk for populations starting below or at the Allee threshold, but a lower extinction risk for populations starting above the threshold (Fig. 2F). The effect takes place over a narrower range of population sizes (Fig. 1), therefore, a population that starts at 75 individuals experiences the full effect and is likely to decline more rapidly (Fig. 2F) than a similar population with a gradual Allee effect (Fig. 2E). In contrast, a population that starts at 125 individuals is above the level where the Allee effect takes place, and therefore will likely increase and recover, whereas with the gradual effect a population starting at 125 individuals had some chance of falling below the threshold. The effect of HCM is similar to the gradual Allee scenario for populations starting below the threshold; the population goes extinct faster. The greatest difference from the gradual Allee scenario occurs for populations starting above the threshold with HCM. With the steep Allee effect and HCM equal to 0.02, the population has a low probability of extinction (0.05) because it is above the population level where the Allee effect decreases the expected growth rate. In contrast, in the gradual case a population starting at 125 individuals with HCM equal to 0.02 had a high probability (0.055) of declining (Fig. 2E,F).

### Combination A

Combining the three factors above at their first setting produces extinction risk without HCM for all initial population levels considered here. Extinction risk was higher for all population size and HCM levels (Fig. 2G) relative to the base Allee effect alone (Fig. 2E). In combination with an Allee effect, catastrophes substantially increased extinction probabilities. For example, for a population starting at 150 individuals with 0.01 HCM, extinction risk was increased 0.20 by adding the other base effects. For populations starting below the Allee threshold, extinction risk is higher because extinction occurs more rapidly when the other factors are added (Figs. 2E,G). For populations starting above the threshold with no or low HCM, the absolute extinction risk is increased because catastrophes or environmental variance can cause the population to fall below the Allee threshold. These factors therefore increase the influence of the Allee effect. In particular, populations above the Allee threshold are no longer buffered from additional extinction risk at low rates of HCM. Extinction risk rises immediately for even the lowest levels of HCM (Fig. 2G).

### Combination B

As expected, the risk of extinction is greater with the combination of the greater effects, even for populations without HCM. The risk is at least 0.45 for all scenarios examined (Fig. 2H). As with the combination of base effects, the risk of extinction rises immediately with HCM, even at the lowest levels, and rises in a nearly linear fashion. Extinction risk is much greater for the populations starting above the Allee threshold than it was with just the greater Allee effect alone (Figs. 2F,H). For example, a population starting at 150 individuals with just the Allee effect and with 0.03 HCM had an extinction risk of 0.06, but that risk was raised to 0.83 by the other factors. Another difference is seen with a population starting at 75 individuals with no HCM, where the correlated environmental variability allows a small chance for the population to escape to levels above the influence of the Allee effect if it has a run of “good” years, whereas in contrast a similar population with just the Allee effect is apparently doomed to extinction. The risk of extinction in 100 years was still greater for the population with the combination of all the greater effects, but there is a slightly greater likelihood of recovering to higher population levels. A population starting at 75 individuals with 0.02 HCM appears to have no chance of escaping extinction. A population starting at 125 individuals with no HCM has an uncertain fate – it might escape and recover or it might (through “bad” years or a catastrophe) fall below the Allee threshold and decline to extinction. The addition of 0.02 HCM makes it highly likely that the population will decline below the threshold and continue to decline to extinction.

Threshold levels of HCM that meet the specified criteria of causing extinction risk to increase by 0.1 over the no-HCM risk level are plotted in Figure 4. HCM would need to average less than 0.33 or 0.5 of an individual per year for populations starting at 100 or less (Figure 4A). For populations starting at 150, HCM would need to be less than 1.5 individuals per year to meet the criteria. Overall there is (as expected) a general trend upwards with population size. When expressed as a percent of the population (4B), a level of HCM of 0.5% would meet the criteria for all scenarios and all starting population sizes).

**Figure 4.**
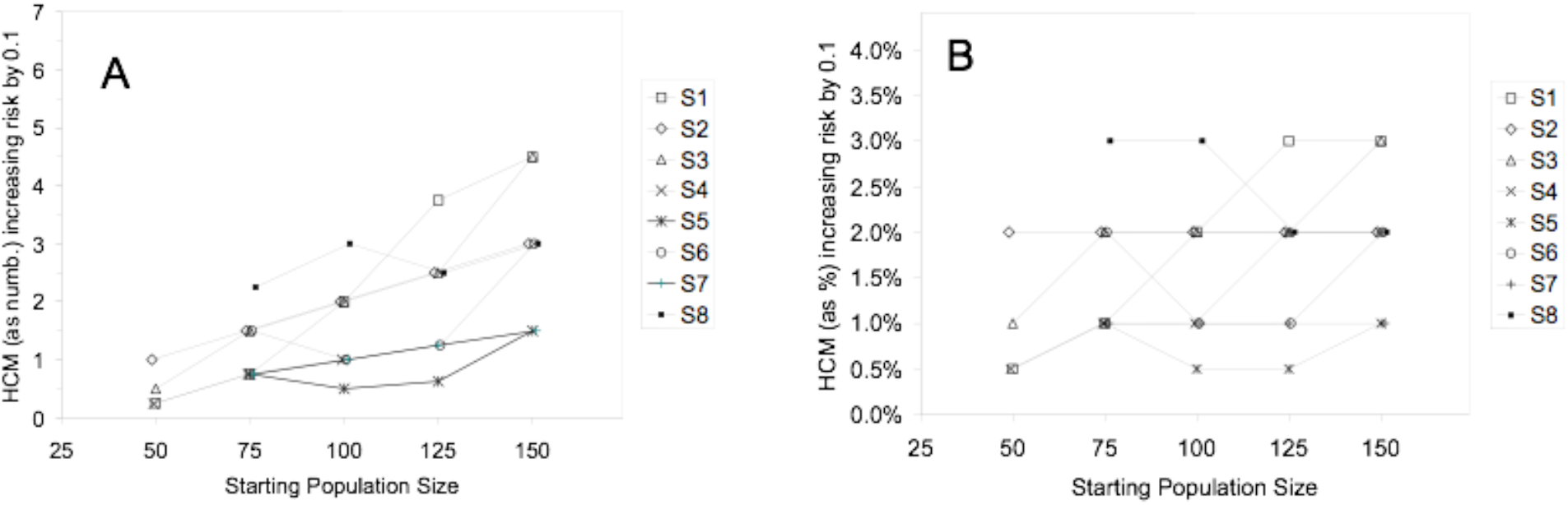
Levels of HCM that increased the absolute risk of extinction by at least 10% compared to a scenario with no HCM, plotted against starting population size, for the eight scenarios in Table 2. The level of HCM is expressed as a number (A) or percentage of the population (B).

## DISCUSSION

Different scenarios of population dynamics led to qualitatively different outcomes for extinction risk, even in the absence of HCM. These results were not unexpected and were consistent with previous examinations of factors affecting extinction risk (e.g., Dennis 1989, Lande 1993). What is important here is to examine how HCM affects extinction risk, and to suggest a method of establishing an acceptable thereshold for HCM of small populations of marine mammals.

### Risk of extinction from different population dynamics models

With no HCM, there was little risk of extinction from un-correlated environmental variance at the level we specified. Even though these populations start at a small initial size (as low as 50 individuals), they are “rescued” by the model’s expected positive population growth, a consequence of the use of a density-dependent model with compensation. The importance of this rescue effect has been described before (Mills *et al.* 1996). The auto-correlated environmental variance we specified led to a greater probability of extinction. Catastrophes had very little effect on extinction risk as a single component within the range explored here. Even with a drop in survival by 0.40 at an expected frequency of once every 33 years, the simulated populations were still “rescued” by the model’s expected positive population growth rate in intervening years. Extinction risk will obviously increase at some greater frequency and/or magnitude of catastrophic events than was tested here. Catastrophes did have an important interaction when considered in combination with the other model scenarios specified here (discussed below). In our range of scenarios the Allee effect had a stronger influence on extinction risk. Compared to the gradual Allee effect (Z_d_ = 1.5), the steeper effect (Z_d_ = 10.0) led to a higher extinction risk for populations starting below or at the Allee threshold but a lower extinction risk for populations starting above the threshold. This is because in our model the steeper Allee effect reduces population growth more rapidly but the effect takes place over a narrower range of population sizes. Our results agreed with those of Dennis (2002), who concluded that Allee effects produce a qualitatively different result for extinction risk than does demographic variance alone (i.e., Allee effects produce a population threshold across which extinction risk changes dramatically). Environmental variance and catastrophes in combination with the Allee effect produced a greater risk of extinction than seen for the Allee effect alone. In other words, the increased risk of extinction was not additive across the three components – there was an interaction between them that increased the risk of extinction. In particular, catastrophes could drop a population from above the Allee threshold to below the Allee threshold where it would face near certain extinction.

### Risk of extinction from HCM

We summarize the effects of HCM on small populations by describing four categories:

1. HCM delays the rate of recovery of a population but does not increase extinction risk.
2. HCM causes a population that would otherwise likely recover to linger at low population levels, increasing its risk of extinction from stochastic processes.
3. HCM causes a population that would otherwise likely recover to have a greater probability of falling below an Allee threshold (if there is one), and thus decline to extinction (if HCM is large enough the probability becomes a certainty).
4. HCM accelerates the rate of extinction for a population that is already doomed to extinction, if current conditions continue.

In scenarios where there was no risk of extinction in the absence of HCM, the addition of HCM did not lead to extinction risk until the level was at the upper range examined here. In other words, for populations that were likely to be “rescued” by strong positive population growth, it took a HCM high enough to counteract that population growth before the risk of extinction would arise. The effect of lower levels of HCM would be to delay the rate of recovery of the population.

In nearly all scenarios where there was some risk of extinction in the absence of HCM, even the lowest levels of HCM examined increased the risk of extinction. For example, for a HCM level of 0.0025 (1/4 of a percent of the population, on average, each year), extinction risk was increased by 0.09 or 31% (0.09/0.29) for the steep Allee scenario with a starting population size of 100, a substantial increase from what is initially just one expected death from HCM every four years. Similarly, for Combination A, for a population starting at 75 individuals the risk of extinction increased from 0.57 to 0.62 with an increase of HCM from zero to 0.0025.

In all the scenarios with an Allee effect where the population started above the Allee threshold, the effect of HCM was to increase the possibility that the population would fall below the threshold and therefore incur increased risk of extinction. The amount of HCM required to cause this effect was a function of how far above the Allee threshold the population started.

In scenarios where the population started below the Allee threshold, HCM had the effect of increasing the rate of extinction for populations that were essentially “doomed”. As expected, populations that started well below the Allee threshold were unable to recover, and any HCM simply hastened extinction. Although the probability of extinction was not always close to 1.0 after 100 years in the simulation, nearly all these trajectories exhibited a decline to very low population sizes and would likely go extinct within the next 50 years or so.

Overall, the threshold results suggest that HCM of less than 0.5% per year would increase the risk of extinction by no more than 10% in most cases. This represents an average of one death every two years for a population size of 100, and one death every four years for a population size of 50. The threshold rose to an HCM 1% or higher for populations starting at a population size of 150 (on average three deaths every two years). It should be noted that incorporation of the Allee effect with the Allee parameter T set at a population size of 100 clearly influenced much of the results for populations starting at 100 or less. However, some scenarios without the Allee effect had similar results, so these results were not only because of the specified Allee effect.

A few caveats are important before considering how general these results might be. The results will be different for different criteria for increase in extinction risk, compared to the 10% increase in extinction risk considered here. Additionally, all scenarios were run with a maximum population growth rate of 4% per year. Although this may be a good default value in the absence of information (see Wade 1998), some species (e.g., humpback whales) can clearly grow at faster rates than this, whereas other species (e.g, sperm whales) may not be able to grow this fast. For populations with species-specific population growth rate information available, these simulations could be repeated in a case-specific way to examine results relative to different maximum population growth rates.

It would also be worth further investigation to determine if a threshold could be found that was a function of both population size and maximum growth rate. In this case, an HCM threshold of 0.5 % of the population size represents 1/8 of the specified maximum growth rate. In the PBR approach to managing bycatch, it was found that a value of ½ the maximum growth rate multiplied by a minimum population size estimate (20^th^ percentile) was found to meet the goal of maintaining at or recovering populations to above the maximum net productivity level (Wade 1998). Further, a value of ¼ the maximum growth rate was suggested to account for potential biases in the data. We found that an even lower value (1/8 of the maximum growth rate) was needed to avoid substantially increasing the extinction risk of a small population. If the value found here was similarly cut in half to account for potential biases to 1/16 of the maximum population growth rate, this would result in a threshold nearly identical to that resulting from a PBR calculated with a “recovery factor” of 0.1 (Wade 1998). This is, coincidentally, the mortality threshold suggested by Wade (1998) based on the criteria of not delaying recovery to the maximum net productivity level by more than 10%.

### Managing HCM for small populations

For small marine mammal populations suspected to have dynamics that include environmental variability, catastrophes and an Allee effect, it is clearly important to avoid HCM or control it at very low levels. Environmental variance in survival or fecundity is likely common in marine mammals, although relatively few datasets are available to demonstrate it. Long-term studies have shown that estimates of variance of population size continue to increase for the first 20 years of data collection, suggesting that long-term studies are needed to capture the effect of severe mortality events (Pimm and Redfearn 1988, Arino and Pimm 1995). Relatively infrequent catastrophic mortality can be viewed as an extreme form of environmental variance (Lande 1993). Catastrophic mortality events have been documented in marine mammals, particularly from disease and biotoxins (Gulland et al. 2002, 2005) and from oceanographic events such as El Nino (Limberger 1990).

Studying the demography of very small populations is difficult, but several studies have indicated that Allee effects do occur in a variety of species (Groom 1998, Kuussaari *et al.* 1998, Courchamp *et al.* 1999), though one meta-analysis study found little evidence for depensation in fish species (Myers *et al.* 1995). Fowler and Baker (1991) concluded that depensation was likely to be a common phenomenon in animal population dynamics, especially in populations smaller than 10% of their original, pre-exploitation populations (0.1K; Pimm and Redfearn 1988, Arino and Pimm 1995). Depensatory effects are clearly an important consideration for assessing the risk of extinction (Fowler and Baker 1991, Dennis 2002), and our results have shown that HCM can particularly affect extinction risk for populations near or below the threshold level for the Allee effect.

In general, we recommend that managers consider the range of potential population dynamics for the species under consideration and make precautionary decisions about the level of HCM that is likely to be sustainable. We currently have little ability to fully understand the population dynamics at low levels for many or most marine mammal populations, and so we will not often be able to unambiguously determine a population’s risk. For critically depleted populations (e.g., at low levels relative to historic levels and at small population sizes), a very small amount of HCM could substantially increase the risk of extinction.

For example, for very small populations like the North Atlantic right whale (Kraus 1990) and some Hector’s dolphin populations (Martien *et al.* 1999, Dawson *et al.* 2001, Burkhart and Slooten 2003), the risk of extinction could be substantially elevated by the human-caused mortality of a small number of animals. For both of these species there has been considerable discussion among scientists and managers on the issue of setting a sustainable level of HCM. In both cases managers have decided that currently available methods of setting allowable levels of HCM (e.g., Wade 1998) are insufficiently precautionary and fisheries bycatch should be reduced as close to zero as is practicable (e.g. Waring *et al.* 2002, Suisted and Neale 2004, DOC and MFish 2006). The methods outlined in this paper make it possible to provide specific management advice in these sorts of cases.

The conundrum for managers is that we have shown that the effect of HCM is conditional on the assumed population dynamics scenario, and consequently the amount of extinction risk a population faces in the absence of HCM. However, for species for which the population dynamics scenarios specified here are thought plausible, managers need to account for this uncertainty when considering the potential effect of HCM. For small populations which are thought to be at risk of extinction (i.e., where there is a risk of extinction due to small population size and/or potential dynamics), one approach would be to focus on simulations of population scenarios that have extinction risk in the absence of HCM, and examine the increased risk of extinction caused by adding HCM. Managers can examine the increase in extinction risk that HCM is predicted to cause, and consider how much additional risk can be tolerated for the human activities that cause the mortality.

## ACKNOWLEDGEMENTS

John Brandon provided substantial help in programming and running the simulations, and we greatly appreciate his efforts. Marcia Muto provided excellent editorial help, as did Jim Lee and Gary Duker. We are grateful to Jay Barlow, Steve Dawson, Rod Hobbs, and Barbara Taylor for formal and informal reviews and discussions of the work in this paper. We particularly thank Ari Samaranayaka and David Fletcher for discussions and for help on error-trapping algorithms regarding sampling from Beta distributions. We also thank Michael Runge for a thorough and helpful review, and for taking the time to check all the equations.

